# Successful spread of *mcr-1*-bearing IncX4 plasmids is associated with variant in replication protein of IncX4 plasmids

**DOI:** 10.1101/2023.11.23.568506

**Authors:** Lingxian Yi, Kaiyang Yu, Guolong Gao, Rongmin Zhang, Luchao Lv, Daojin Yu, Jun Yang, Jian-Hua Liu

## Abstract

IncX4 plasmids are one of the most epidemiologically successful vehicles for *mcr-1* spread. Here we found that the IncX4 plasmids carried two different replication proteins encoded by genes *pir-1* and *pir-2*, respectively, but *mcr-1* was only carried by IncX4 plasmid encoding *pir-1*. The copy number of *pir-2* encoding plasmids (3.15±0.9 copies) are higher than that of *pir-1* encoding plasmids (0.85±0.5 copies). When *mcr-1* was cloned into IncX4 plasmid encoding *pir-2*, the higher copy number of these plasmids resulted in increased expression of *mcr-1* and a greater fitness burden on their host cells. However, these plasmids exhibited a lower rate of invasion into the bacterial population compared to *mcr-1* positive plasmids encoding the *pir-1* gene. These findings collectively explain the absence of *mcr-1* in all IncX4 plasmids encoding *pir-2*. Our results further confirmed that low-copy numbers are important for the spread of *mcr-1* plasmid from the perspective of natural evolution.

## Introduction

Plasmid, as extrachromosomal DNA, is capable of replicating and transferring in different bacteria and usually carries genes which help hosts adapt to diverse environments, such as antibiotic exposure^1^. Based on the merit, plasmids become the perfect vectors for the dissemination of antibiotic-resistant genes^2^. Plasmid transfer is also a major driving force for the rapid spread of colistin-resistant gene, *mcr-1*^3^. Up to date, *mcr-1* gene has spread in samples of diverse origins in over 61 countries and regions^4^, while the global transfer of IncX4 and IncI2 plasmids carrying *mcr-1* is the major reason for the *mcr-1* outbreak^5^. The epidemic success of IncX4 plasmids is strongly correlated with *mcr-1* carriage. After withdrawing colistin as an animal food addictive in China, the detection of IncX4 plasmids was dramatically reduced^6^. In addition, IncX4 plasmids usually carry a single antibiotic-resistant gene, mostly *mcr-1*. IncX4 plasmids belong to the IncX family. By far, the IncX family plasmids have been grouped into 9 subgroups based on their resolvase proteins, and most of them are common vectors for clinically important antibiotic-resistant genes, for example, the *bla*_NDM_ carriage by IncX3, *tet*(X4) carriage by IncX1, and *mcr-1* carriage by IncX4, while the mechanism involved in such strong relationship between plasmids and antibiotic-resistant genes remains unclear^7,8^. Hence, in this study, we aim to investigate the mechanism involved in the global prevalence of *mcr-1*-bearing IncX4 plasmids. By comparing and analyzing the genetic characterization of 351 complete IncX4 plasmid sequences, we found IncX4 plasmids encoded two homologous replications, which caused an average ∼3.7-fold difference in plasmid copy number and have a significant impact on the persistence of *mcr-1-* bearing IncX4 plasmids.

## Results and Discussion

### Characterization and phylogenetic analysis of IncX4 plasmids

A total of 351 IncX4 plasmid sequences, including 338 plasmid sequences deposited in GenBank and 13 plasmid sequences obtained through illumine sequencing in our lab, were used in this study. These IncX4 plasmids were carried by strains recovered from livestock, animal product, environments, and humans, from China, USA, Brazil, UK, and other countries between the 1980s and 2022, and 232 of the 351 plasmids carried *mcr-1* genes. A total of 192 out of the 351 IncX4 plasmids shared clear information on the collection dates. To better understand the sequence evolution of IncX4 plasmids, we constructed a maximum likelihood tree of the variations in the core genomes of the 351 IncX4 plasmid sequences (Figure 1). We observed a high degree of homology among the 232 *mcr-1*-bearing IncX4 plasmids with an average of 1.12 SNPs±3.90 distance. In contrast, the remaining 119 *mcr-1* negative IncX4 plasmids exhibited an average of 52.13±54.25 SNP differences. The highly conserved construction of the sequences of the 232 *mcr-1*-bearing IncX4 plasmids suggests that all *mcr-1*^+^ IncX4 plasmids probably originated from a common predecessor. Given the minimal SNP difference among these plasmids, it is likely the acquisition of *mcr-1* by IncX4 plasmids took place in recent years, followed by successful global dissemination. The earliest known acquisition of *mcr-1* in IncX4 plasmids so far can be traced back to 2009, however, IncX4 plasmids have emerged in the 1980s (Table S3).

**Figure 1.**
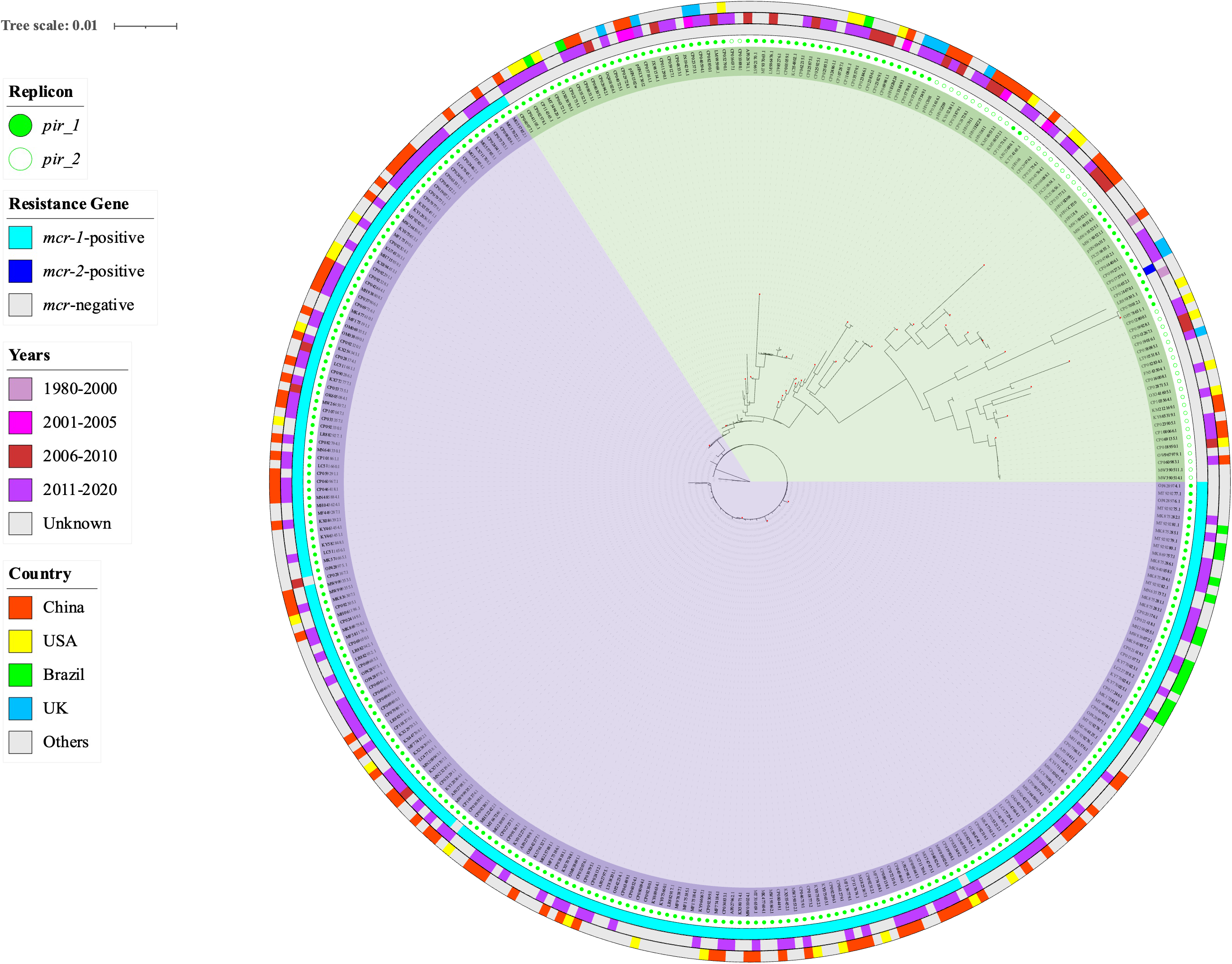
Phylogenetic tree of 351 IncX4 plasmids including 338 IncX4 plasmids deposited in GenBank and 13 IncX4 plasmids in our lab. Purple - The plasmid sequences showed high similarity in the SNP core genome; Green - the plasmid sequence showing less similarity; Light blue rectangles – presence of *mcr-1*; Dark blue rectangles-lack of *mcr-1*; Gray rectangles - presence of *mcr-2*; Filled green circle - presence of *pir-1*; Empty green circle – presence of *pir-2*; The outer and second rings surrounding the phylogenetic tree - the collecting year and countries of IncX4 plasmids; Red dots - the representative plasmids used for linear sequence comparison.

By sequence analysis of the 192 IncX4 plasmids with clear collecting date, we found a variance of antimicrobial resistance gene carriage in the IncX4 plasmids over time. Before 2010, a total of 30 IncX4 plasmid sequences were obtained, only 7(23%) of them carried resistance genes, one plasmid carried *dfrA1* genes, 3 plasmids possessed *bla*_CTX-M-27_, and 3 plasmids carried *mcr-1* (Table S3). However, of 162 IncX4 plasmid sequences obtained after 2010, 112(69.1%) were found to carry *mcr-1* gene, 40 (24.1%) did not carry any resistance genes, and 10 plasmids carried other resistance genes, including *dfrA1*, *qnrS*, *bla*_CTX-M-121_, *bla*_CTX-M-14_, *cfr, fosA, ermB, tet(*M) or *bla*_TEM_. The tendency of *mcr-1* positive plasmids to undergo high-throughput sequencing for further research may have contributed to their predominance among plasmids isolated after 2010.

### Variation in plasmid replication proteins and plasmid copy number

Three *mcr-1* positive plasmids (SNP=1), and 23 *mcr-1* negative plasmid sequences (SNP>12) were randomly selected as representative plasmids for linear comparison of the complete plasmid sequence. The results showed that the whole sequence of *mcr-1* positive plasmids displayed a high similarity with >99.97% identity. However, for the *mcr-1* negative plasmids, though the genes related to the function of plasmid conjugation and DNA processing exhibited high identity, sequence variations were observed, including the diversity in the *pir* gene, antibiotic-resistant genes, and certain genes with unknown functions inserted in the variable region (Figure S1). Two *pir* genes encoding two homologous replication proteins with 28.67% amino acid identity were identified in IncX4 plasmids (Figure S2). Previously, these two homologous genes were both annotated as *pir* genes, they hereafter were renamed as *pir-1* and *pir-2*, respectively, for distinction. Notably, it was observed that the *mcr-1* gene is present on the IncX4 plasmid with *pir-1* gene, but not IncX4 plasmid with *pir-2* gene (Figure 1).

In order to know the evolution history of IncX4 plasmids, Bayesian analysis was performed to trace the earliest emergence of *pir-1* and *pir-2* plasmids. The phylogenetic tree indicated that the ancestral nodes of *pir-2* plasmids emerged in the year 1980 (Figure 2A). A previous study revealed that *mcr-1* has emerged in *E. coli* isolates derived from the 1980s when *pir-2* plasmids already existed^9^. But why there is no *mcr-1* gene in *pir-2* encoding IncX4 plasmids?

**Figure 2.**
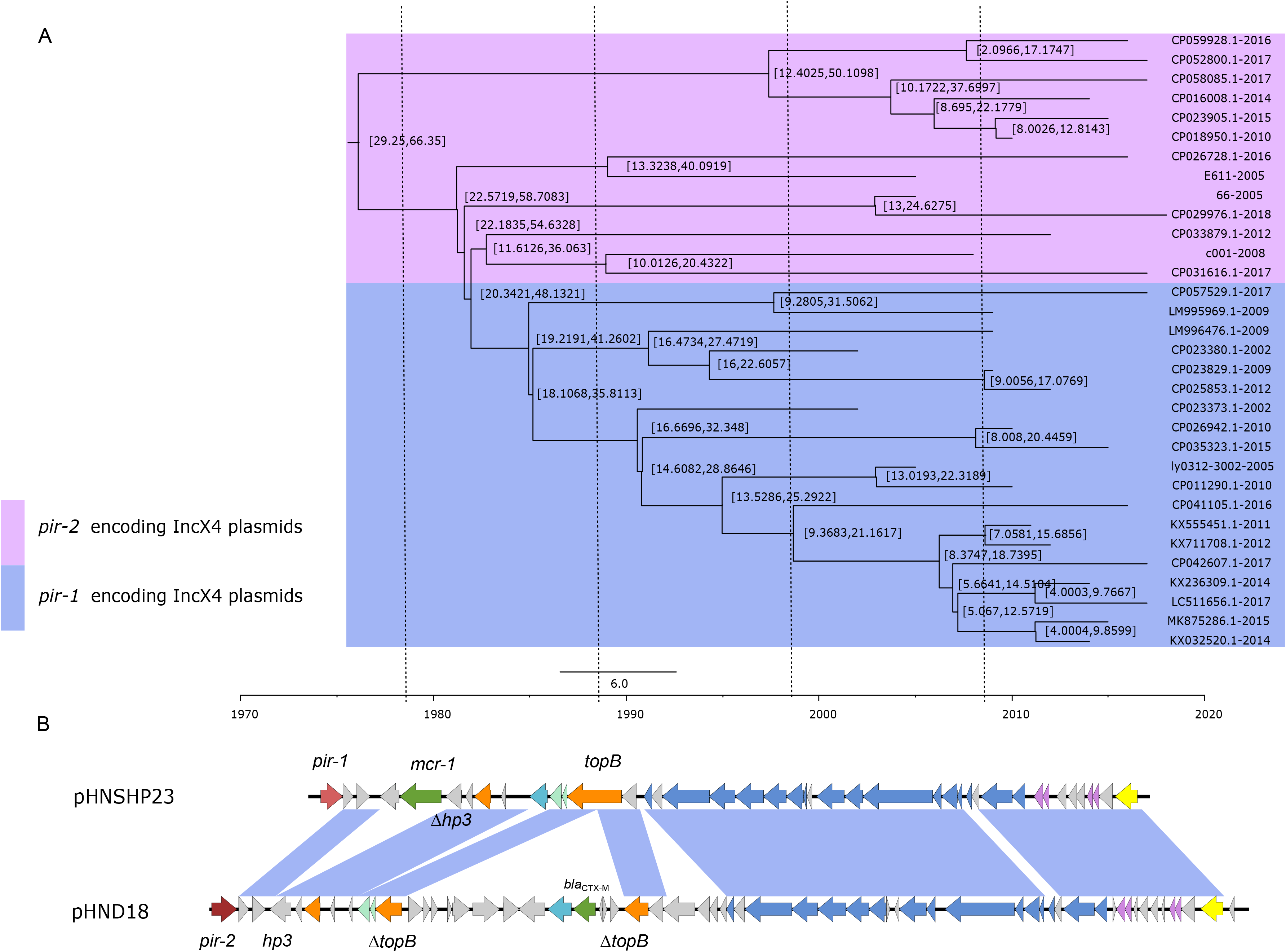
Bayesian analysis and sequence comparison of representative IncX4 plasmids with *pir-1* and *pir-2*. Blue – 19 *pir-1* encoding IncX4 plasmids; Purple - 13 *pir-2* encoding IncX4 plasmids. Timescale is displayed at the bottom. B. Sequence comparison of pHNSHP23 (*pir-1*) and pHND18(*pir-2*) plasmids. Genes are labeled in different colors based on their function. Red - replication genes; Green - antibiotic-resistant genes; Light blue - mobile elements; Blue – genes related to plasmid conjugation; Purple - toxin-antitoxin systems encoding genes; Gray - genes with unknown function; Orange - genes related to DNA processing.

The process of plasmid replication was tightly controlled by plasmid replication protein^10^. Mutations in replication genes frequently caused a change of plasmid copy number^11^. Our previous study revealed that the increased copy number of *mcr-1*-bearing IncI2 plasmids imposes a higher fitness burden on the host, resulting in reduced persistence of the plasmids in bacteria^12^. Given the significant correlation between *pir* gene sequences and *mcr-1* persistence in IncX4 palsmids, we hypothesize that the *pir* gene plays a crucial role in facilitating the successful dissemination of the IncX4 plasmids carrying *mcr-1* genes. To confirm the hypothesis, the copy number of 14 IncX4 plasmids, including 5 *pir-1* encoding and 9 *pir-2* encoding plasmids, were then precisely determined by qPCR. The copy number of *pir-1* encoding plasmids was estimated to be 0.85±0.5 copies per chromosome, while 3.15±0.9 copies of *pir-2* encoding plasmids in wild-type strains, indicating that the copy number of *pir-2* encoding plasmids are higher than that of *pir-1* encoding plasmids (Figure 3A).

**Figure 3.**
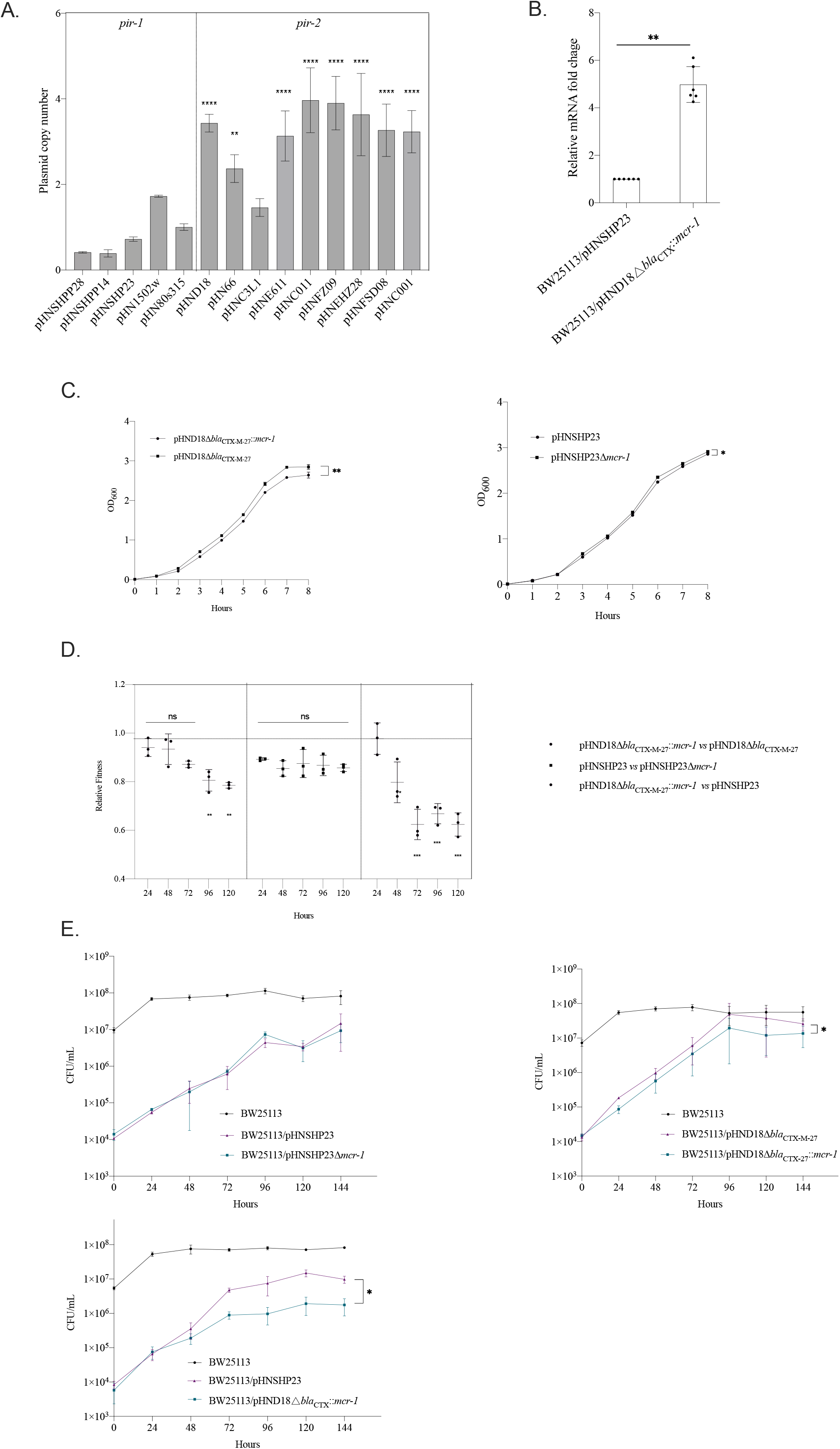
Variants in plasmid copy numbers, mRNA level of *mcr-1*, growth kinetics, host fitness, and invasion capability caused by two homologues of replication protein. A. Estimation of plasmids copy number. The copy number of each plasmid was measured by qPCR and Illumina sequencing using the *gapdh* gene as a reference. The bar represents the mean and SD of biological triplicates. The statistic analysis was performed by a one-way ANOVA with Dunett’s comparison; B. The mRNA level of *mcr-1* in BW25113/pHNSHP23 and BW25113/pHND18Δ*bla*_CTX-M-27_::*mcr-1*. The data was assessed by two-tailed paired *t*-test with Wilcoxonsigned rank test. C. Growth curves of *E. coli* BW25113 carrying plasmid pHNSHP23, pHND18, and their Δ*mcr-1* mutants. The data was assessed by two-tailed paired *t*-test with Wilcoxon signed rank test. D. *E.coli* BW25113/pHNSHP23 was competed with BW25113/pHNSHP23Δ*mcr-1* or BW25113/pHND18Δ*bla*_CTX-M-27_::*mcr-1* in *vitro*, respectively. BW25113/pHND18Δ*bla*_CTX-M-27_ was competed with BW25113/pHND18Δ*bla*_CTX-M-27_::*mcr-1* in *vitro*. The Y-axis Relative fitness indicated: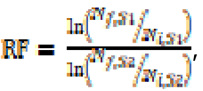 where RF is the related fitness of strain S1 compared to strain S2 and N_f,sx_ and N_i,sx_ correspond to the CFU counts of strain Sx at the observed timepoint and at the initial timepoint, respectively. All competition assays were performed with three biological replicates. a One-way ANOVA with Dunnett’s comparison test were performed on the RF value of each bar compared to the RF value of 24h. The data was assessed by two tailed *t*-test. E. Co-cultures of BW25113/pHNSHP23Δ*mcr-1* or BW25113/pHND18Δ*bla*_CTX-M-27_::*mcr-1* with BW25113/pHNSHP23 and 1 000-fold excess of BW25113 without any plasmids. Co-cultures of BW25113/pHND18Δ*bla*_CTX-M-27_::*mcr-1* with BW25113/pHND18Δ*bla*_CTX-M-27_ and plasmid-free BW25113. *P*-value was calculated by a two-tailed paired *t*-test with Wilcoxon signed rank test. Statistical significance is indicated as follows:****, P<0.0001; ***, P<0.001; **, P<0.01; *, P<0.05; ns, not significant.

### Evaluation of the persistence of mcr-1-bearing pir-1 or pir-2 encoding IncX4 plasmids

It is known that high-expression of *mcr-1* is toxic for host cell and impairs its host cell growth, thus the copy numbers of *mcr-1* positive plasmids are commonly very low^12,13^. Considering the copy number difference among *pir-1* and *pir-2* encoding IncX4 plasmids, we suspected that the *mcr-1* expression in *pir-2* plasmids would be higher than *pir-1* plasmids due to the copy number increase. To confirm the suspect, the plasmids pHNSHP23 and pHND18 were chosen as the representatives for IncX4 plasmids carrying *pir-1* and *pir-2* genes, respectively. Sequence comparison of pHNSHP23 and pHND18 confirmed the conserved backbone sequence and the acquisition of different resistance genes. The loci of resistant genes in these two plasmids were also different, the *mcr-1*-*pap2* cassette was inserted within a hypothetic gene *cds3* of pHNSHP23, whereas a genetic cassette *orf*s-IS-*bla*_CTX-M-27_-*orf*s was inserted in the *topB* gene of pHND18 plasmid backbone (Figure 2B). The genetic cassette including *bla*_CTX-M-27_ was replaced by *mcr-1* through the λRed homologous recombination method to obtain mutant pHND18△*bla*_CTX-M-27_::*mcr-1*. We then identified the expression of *mcr-1* in pHNSHP23 and pHND18△*bla*_CTX-M-27_::*mcr-1* plasmids by RT-qPCR. The result showed that the expression of *mcr-1* in pHND18△*bla*_CTX-M-27_::*mcr-1* is ∼4.98-fold higher than that in pHNSHP23 (Figure 3B).

Our precious results indicated that a high expression level of *mcr-1* as a result of increased plasmid copy number would contribute to a high fitness cost on its hosts. Hence, we suspected that the higher expression of *mcr-1* in *pir-2* plasmids affects their persistence in the bacterial population, resulting in the absence of *pir-2* encoding IncX4 plasmids carrying *mcr-1* gene. We compared bacterial fitness of *mcr-1* carriage by these two types of IncX4 plasmids with different plasmid copy numbers. Firstly, we tested the impact of *mcr-1* carriage on the host growth kinetics by pHNSHP23 or pHND18, respectively. The results showed that compared to pHND18△*bla*_CTX-M-27,_ pHND18△*bla*_CTX-M-27_::*mcr-1* carriage caused a slight reduction in strains growth during 8 h evaluation, while no difference was observed between pHNHP23 and pHNSHP23△*mcr-1*(Figure 3C). Competition assays between pHNHP23 and its mutant pHNSHP23△*mcr-1*, mutants pHND18△*bla*_CTX-M-27_ and pHND18△*bla*_CTX-M-27_::*mcr-1*, and pHNSHP23 and pHND18△*bla*_CTX-M-27_::*mcr-1* were carried out. We observed that *mcr-1* carriage by IncX4 plasmids at high copies or low copies both caused a fitness burden to *E. coli* host BW25113 (Figure 3D). However, the fitness caused by high-copy IncX4 plasmids was more obvious than that by low-copy IncX4 plasmids. Compared to △*mcr-1* mutant plasmids pHND18△*bla*_CTX-M-27_ or pHNSHP23△*mcr-1*, the relative fitness of pHND18△*bla*_CTX-M-27_::*mcr-1* reduced dramatically over 5 days, decreasing from 0.94±0.04 to 0.79±0.01, while pHNSHP23 displays a slight reduced relative fitness, from 0.89±0.01 to 0.86±0.02. In competition co-cultures of pHNSHP23 and pHND18△*bla*_CTX-M-27_::*mcr-1*, pHNSHP23 conferred an obvious fitness advantage to *E.coli* (Figure 3D). Collectively, these data suggest that the fitness cost incurred by the *mcr-1* gene on IncX4 plasmids at low-copy number is lower compared to IncX4 plasmids at high-copy number. We also found that even a single copy *mcr-1* carried by IncX4 plasmids at low copy number imposed a fitness burden on its host bacterium and negatively affected the persistence of the *mcr-1*-bearing IncX4 plasmid in the bacterial population. These results are consistent with our previous study on IncI2 plasmid carrying *mcr-1* and help explain the decline of *mcr-1* positive plasmids, specifically IncI2 and IncX4 plasmids, following the ban on the use of colistin as growth promoter^6,12^. However, the underlying molecular mechanism of fitness cost caused by *mcr-1* remain poorly understood and require further exploration.

We tested the capability of WT and mutant plasmids to invade plasmid-free cells. We found that though the conjugative frequency of pHND18△*bla*_CTX-M-27_::*mcr-1* is similar to pHNSHP23 (4.37×10^-4^ *vs* 3.16×10^-4^), pHNSHP23 invaded cells at a much higher rate than pHND18△*bla*_CTX-M-27_::*mcr-1* after 24 h in competition cultures. In addition, pHNSHP23 and pHNSHP23△*mcr-1* showed a mild difference in invasion capability after 96 h, while pHND18△*bla*_CTX-M-27_ invaded cells quicker than pHND18△*bla*_CTX-M-27_::*mcr-1* after 24 h(Figure 3E). These results indicated that the plasmid copy number has an impact on the invasion capability of *mcr-1* IncX4 plasmids, just like *mcr-1*-bearing IncI2 plasmids^12^. Worthy of note, stability of pHNSHP23 and pHND18Δ*bla*_CTX-M-27_ remained unaffected by the *mcr-1* deletion or replacement over 6 days in pure culture(Figure S3).

In conclusion, the fitness cost and the deficiency of invasion capability caused by high copy *mcr-1* bearing IncX4 plasmids explains the lack of *mcr-1* in all *pir-2* encoding IncX4 plasmids. From the perspective of natural evolution, our results further confirmed that low copy number control plays a key role in *mcr-1* plasmid persistence.

## Materials and Methods

### Bacterial isolates and sequencing

Information on strains and plasmids used in this study was listed in Table S1. A total of 13 strains carrying IncX4 plasmids collected from animals in China were included (Table S3). Primers IncX4-F and IncX4-R were used to screen strains carrying IncX4 plasmids (Table S2). All strains were subjected to whole-genome sequencing (WGS) using Illumina HiSeq platforms (San Diego, CA, USA). Illumina reads assemblies were generated using Unicycler version 0.4.3.8. The contigs containing IncX4 plasmid sequence were extracted by PlasmidFinder (https://bitbucket.org/genomicepidemiology/plasmidfinder_db/src/master/). The gaps between contigs were linked by PCR and Sanger sequencing. Plasmids were annotated and analyzed by using the RAST server (https://rast.nmpdr.org), ResFinder (https://cge.cbs.dtu.dk/services/ResFinder/), BLAST (https://blast.ncbi.nlm.nih.gov/Blast.cgi), and SnapGene version 4.3.6.

### Phylogenomic analysis and Bayesian analysis of IncX4 plasmid sequences

Date to May 2023, 338 complete IncX4 plasmids were downloaded from the NCBI database by using the *rep* sequence of IncX4 plasmids extracted from the PlasmidFinder database. Panaroo and RAxML were used to obtain the core genome of 351 IncX4 plasmids and construct a phylogenomic tree^14,15^. SNPs were identified by running Snippy (https://github.com/tseemann/snippy) on the core genome alignment. SNPs compared to the reference plasmid sequence pHNSHP23 were counted from the core SNP alignment with snp-dists 0.6.3 (https://github.com/tseemann/snp-dists). Bayesian analysis of molecular sequences was performed according to the previous study^16^. Briefly, 32 representative IncX4 plasmids of each branch in the phylogenomic tree were selected to generate a chronogram using Bayesian evolutionary analysis version 1.10^17^. Tracer v1.7.1 was used to assess convergence using all parameter effective sampling sizes 21 of > 200^18^. LogCombiner v2.6.1 was used to combine tree files and a maximum clade credibility tree was created using TreeAnnotator v2.6.0^18^. iTOL (https://itol.embl.de) and Figtree version 1.4.2 were used to visualize and annotate phylogenetic trees. Protein sequences were aligned using MUSCLE 3.8.31. Gene organization diagrams were generated with Easyfig 2.2.2.

### Construction of plasmids

Deletion mutants were constructed in *E. coli* BW25113 via homologous recombination following the previously described method^12^. Briefly, the plasmids pHND18Δ*bla*_CTX-M-27_, pHND18Δ*bla*_CTX-M-27_::*mcr-1*, and pHNSHP23Δ*mcr-1* were obtained by using primers ctx-d-F/R, ctx-mcr-F/R and mcr-d-F/R, respectively. Primers ctx-c-F/R and mcr-c-F/R were used for confirming the deletion mutants. pKD4 was used as template for PCR amplification. Plasmid pCP20 was used for removing Kanamycin resistant marker.

### Determination of plasmid copy number, plasmid conjugation frequency, invasion capability, and bacterial fitness in vitro

The details methods involved in determination of plasmid conjugation frequency, invasion capability, and bacterial fitness in *vitro* are described in Text S1.

### Statistical analysis and figures

Prism 9 (GraphPad Software) was used to generate graphics and perform statistical analyses. Figures were prepared with Inkscape 1.0.1 (https://inkscape.org/).

## Acknowledgements

This work was supported by the National Natural Science Foundation of China (grants no. 31830099 and 32141002), the National Key Research and Development Program of China (no. 2022YFC2303900), and the Laboratory of Lingnan Modern Agriculture Project (NT2021006).

## Conflict of interest

The authors declare no competing interests.

## Notes

### Competing Interest Statement

The authors have declared no competing interest.

