## Supplementary material for "Successful spread of *mcr-1*-bearing IncX4 plasmids is associated with variant in replication protein of IncX4 plasmids"

**LEGENDS FOR SUPPLEMENTARY MATERIAL**

Text S1. The methods involved in determination of plasmid copy numbe, plasmid conjugation frequency, invaded capability, and bacterial fitness *in vitro.*(DOCX)

Table S1. The information of strains and plasmids used in this study. (DOCX)

Table S2. Oligonucleotides used in this study. (DOCX)

Table S3. Characteristics of IncX4 plasmids in Genbank database and our lab. (XLSX)

Figure S1. Comparative analysis of 26 IncX4 representative plasmid sequence. (DOCX)

Figure S2. The comparison of amino acid sequence of PiR-1 and PiR-2 protein. (DOCX)

Figure S3. In vitro stability of pHNSHP23, pHNSHP23∆*m**cr-1*, pHND18∆*bla*_CTX-M-27_ and pHND18∆*bla*_CTX-M-27_::*mcr-1* in *E.coli* BW25113.

**Text S1. Materials and methods**

**Determination of** **Copy Number via qPCR**

To date, a total of 13 strains carrying IncX4 plasmids, which were isolated before 2010, have been identified in our lab. Among these strains, nine (C011, D18, 66, E611, C3L1，FZ09, EHZ28, FSD08 and C001) encode *pir-2* genes, while four stains (1502w, 80s315, EGM24, LY3012) encode *pir-1* genes. We didn’t find any *pir-2*-encoding IncX4 plasmids within the strains isolated after 2010. Hence, we used these 9 *pir-2*-encoding strains for plasmids copy number analysis. Regarding *pir-1* encoding strains, we randomly selected 2 (1502w and 80s315) from those isolated before 2010, and 3 strains (SHPP28, SHPP14 and SHP23) carrying *pir-1*-encoding IncX4 plasmids collected after 2010. The relative plasmid copy number was measured by qPCR according to previous methods(1). The genomic DNA of *pir-1* encoding stains SHP28, SHP14, SHP23, 80S315,1502w and *pir-2* encoding strains C011, D18, 66, E611, C3L1，FZ09, EHZ28, FSD08 and C001 were extracted using Hipure Bacterial DNA Kit (Magen, China) following the manufacturer’s instructions. The DNA mixture, obtained from all samples, was subjected to a tenfold gradient-diluted and amplified to construction a standard curve. The standard curved was used to calculate the amplification efficiency (E) and correlation coefficient values (R^2^) for the primers. The amplification efficiency (E) for each pair of primers ranges from 90% to 110%, and the correlation coefficient values (R^2^) range from 0.98 to 1. Plasmid gene *pir* and chromosomal gene *gapdh* were selected as targets for qPCR analysis. The plasmid copy number was quantified and determined as the ratio of plasmid DNA to chromosome DNA by the 2^-∆∆Ct^ method.

**Determination of plasmid conjugation frequency, invaded capability, and bacterial fitness in *vitro***

Conjugation experiments were perfomed as described elsewhere by using streptomycin-resistant *E.coli* C600 as recipients(2). The conjugants were selected on LB agar supplemented with colistin (2 μg/mL) and streptomycin (3000 μg/mL). Plasmid conjugative frequency were calculated as the ratio of transconjugats over recipient CFUs. The strain BW25113/pHNSHP23 was used to compete against BW25113/pHNSHP23Δ*mcr-1*, BW25113/pHND18∆*bla*_CTX-M-27_ was used to compete against BW25113/pHND18∆*bla*_CTX-M-27_::*mcr-1* and the strain BW25113/pHND18∆*bla*_CTX-M-27_::*mcr-1* was used to compete against BW25113/pHNSHP23. All competition assays were carried out in biological triplicates following the previous method(3). Briefly, the overnight cultures of the two competitors were mixed at the rate of 1:1. Cultures were 1:1,000 diluted into fresh LB broth every 24 h for 5 days. At each 24 h timepoint, cultures were serially diluted and plated on LB agar plates or LB agar plates containing colistin (Cl). Considering BW25113/pHNSHP23 and BW25113/pHND18∆*bla*_CTX-M-27_::*mcr-1* could both be grown on Cl plates, we distinguished these two strains by colony PCR with primers 23_F/R and D18_F/R (Table S2).

Plasmid invasion assays were performed according to the previous study(3). Briefly, overnight cultures of the recipient strain BW25113 were diluted 1:100 in LB medium and mixed with a 1:10,000 dilution of the overnight cultures of two donor strains (either BW25113/pHNSHP23 and BW25113/pHNSHP23Δ*mcr-1*, or BW25113/pHND18∆*bla*_CTX-M-27_::*mcr-1* and BW25113/pHND18∆*bla*_CTX-M-27_ or BW25113/pHND18∆*bla*_CTX-M-27_::*mcr-1* and BW25113/pHNSHP23Δ*mcr-1*). Cultures were grown in 50 mL tubes containing 2 mL LB at 37 ºC with slow rolling (80 rpm). At each 24 h, cultures were diluted 1:100 into fresh LB broth. Viable counts were gathered at 24, 48, 72, 96, 120, and 144 h by plating serially diluted cultures on non-selective LB agar. All colonies on LB plates were tested by PCR. IncX4-F/R was used to distinguish plasmid-free cells and plasmid-containing cells. BW25113/pHNSHP23 and BW25113/pHNSHP23Δ*mcr-1* or BW25113/pHND18∆*bla*_CTX-M-27_::*mcr-1* and BW25113/pHND18∆*bla*_CTX-M-27_ were distinguished using primer mcr_F/R. BW25113/pHND18∆*bla*_CTX-M-27_::*mcr-1* and BW25113/pHNSHP23Δ*mcr-1* were distinguished by primers 23_F/R and D18_F/R.

Table S1. Strains and plasmids used in this study

| Strain or Plasmid | Relevant genotype or phenotype | Reference |
| --- | --- | --- |
| *Escherichia coli* |  |  |
| BW25113 | F^_^Δ(araD-araB)567 ΔlacZ4787(::rrnB-3) λ^_^rph-1 Δ(rhaD-rhaB)568 hsdR514 | (4) |
| C600 | (Sm^R^) | (5) |
| BW25113/pHNSHP23 | BW25113 harboring the pHNSHP23 plasmid |  |
| BW25113/pHNSHP23Δ*mcr-1* | BW25113 harboring the pHNSHP23Δ*mcr-1* plasmid |  |
| BW25113/pHND18 | BW25113 harboring the pHND18 plasmid |  |
| BW25113/pHND18Δ*bla*_CTX-M-27_ | BW25113 harboring the pHND18Δ*bla*_CTX-M-27_ plasmid |  |
| BW25113/pHND18Δ*bla*_CTX-M-27_:*:mcr-1* | BW25113 harboring the pHND18Δ*bla*_CTX-M-27_::*mcr-1* plasmid |  |
| SHPP28 | Strain harboring *pir-1*-encoding IncX4 plasmids |  |
| SHPP14 | Strain harboring *pir-1*-encoding IncX4 plasmids |  |
| SHP23 | Strain harboring *pir-1*-encoding IncX4 plasmids |  |
| 1502w | Strain harboring *pir-1*-encoding IncX4 plasmids |  |
| 80s315 | Strain harboring *pir-1*-encoding IncX4 plasmids |  |
| D18 | Strain harboring *pir-2*-encoding IncX4 plasmids |  |
| 66 | Strain harboring *pir-2*-encoding IncX4 plasmids |  |
| C3L1 | Strain harboring *pir-2*-encoding IncX4 plasmids |  |
| E611 | Strain harboring *pir-2*-encoding IncX4 plasmids |  |
| C011 | Strain harboring *pir-2*-encoding IncX4 plasmids |  |
| FZ09 | Strain harboring *pir-2*-encoding IncX4 plasmids |  |
| EHZ28 | Strain harboring *pir-2*-encoding IncX4 plasmids |  |
| FSD08 | Strain harboring *pir-2*-encoding IncX4 plasmids |  |
| C001 | Strain harboring *pir-2*-encoding IncX4 plasmids |  |
| Plasmids |  |  |
| pHNSHP23 | The *mcr-1*-bearing IncX4 plasmid with *pir-1* gene |  |
| pHNSHP23Δ*mcr-1* | The pHNSHP23 plasmid with deletion of *mcr-1* gene |  |
| pHND18 | The *bla*_CTX-M-27_-bearing IncX4 plasmid with *pir-2* gene |  |
| pHND18Δ*bla*_CTX-M-27_ | The pHND18 plasmid with deletion of *bla*_CTX-M-27_ gene |  |
| pHND18Δ*bla*_CTX-M-27_:*:mcr-1* | The pHND18 plasmid with substitution of *bla*_CTX-M-27_ gene by mcr-1 gene |  |
| pKD4 | Km^R^, PCR template for one-step chromosomal gene inactivation | (4) |
| pCP20 | Thermo-inducible expression of Flp recombinase (Ts^R^ Ap^R^ Cm^R^) | (6) |
| pKD46 | Ap^R^, λRed recombinase expression | (4) |

Ap, ampicillin; Km, kanamycin, Ts, thermosensitive.

Table S2. Oligonucleotides used in this study.

| Primers | Sequence |
| --- | --- |
| IncX4-F | AGCAAACAGGGAAAGGAGAAGAT |
| IncX4-R | CTGTCGGCATGTCTGTCTC |
| ctx-mcr-F | TGGTAAAGTAAGTTATCCCTGTCGAGATACTGAAAAGCGAGATACTGAAGGTATAACTTTCGATAA |
| ctx-mcr-R | GCGGCTCTGCCGCCAGACGTAATCGCCGGTTAGGTTGATCGAATGGAGTGTGCGGTGGGTTTGGAAAA |
| ctx-d-F | TGGTAAAGTAAGTTATCCCTGTCGAGATACTGAAAAGCGTGTAGGCTGGAGCTGCTTCG |
| ctx-d-R | GCGGCTCTGCCGCCAGACGTAATCGCCGGTTAGGTTGATATGGGAATTAGCCATGGTCC |
| mcr-d-f | GAGAAACTACTCAAAAAATAAACGGTGGGATAATTGCGGTGTAGGCTGGAGCTGCTTCG |
| mcr-d-r | ACGCCCAAGCGATCCAGCGTATCCAGCACATTTTCTTGGTCCATATGAATATCCTCCTT |
| mcr-c-F | GCGACCATCAGATCAATCTGACT |
| mcr-c-R | CGCAATTATCCCACCGTTT |
| ctx-c-f | CAAAAATGATTGAAAGGTGGTTG |
| cxt-d-r | TTGGTGATTTTGAACTTTTGCTT |
| 23_F | GATTGATAGCCGTAAACCAC |
| 23_R | TATCAACAGCAGGATCAAGCA |
| D18_F | CGCTTAACAAAAGTGAGACA |
| D18_R | ACAGATAATAAAACTTGCGTTG |

Figure S1

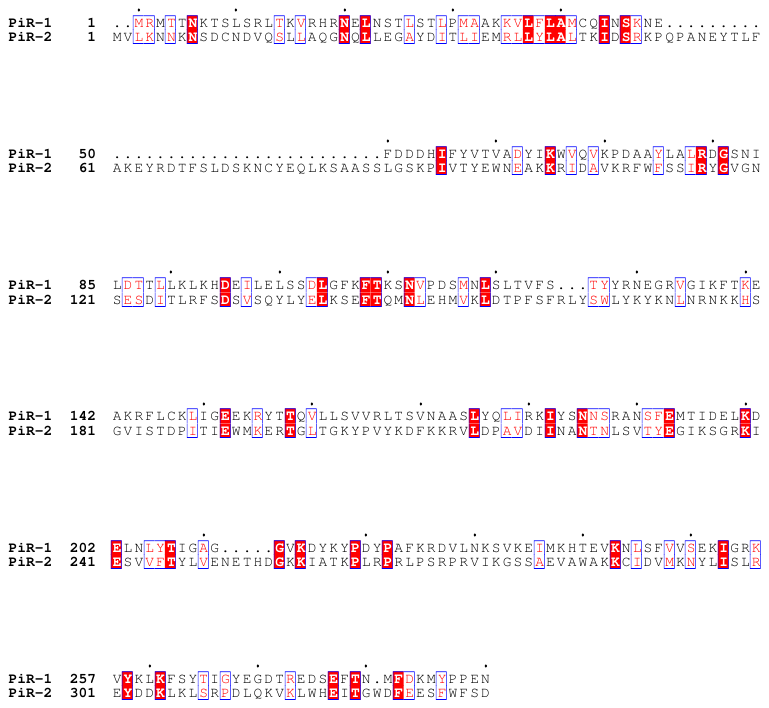
Figure S2

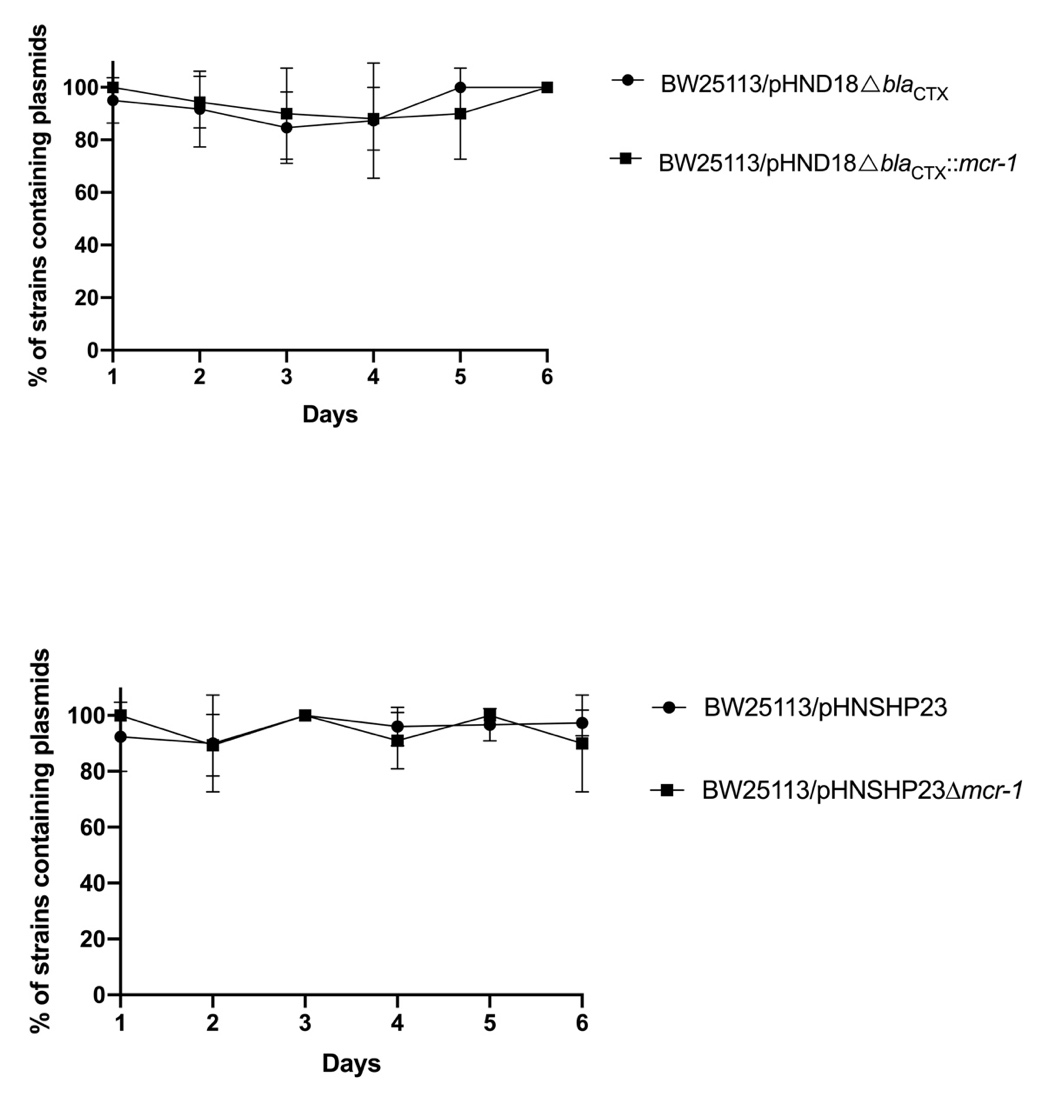
Figure S3
