## Supplementary figures and images for "Successful spread of *mcr-1*-bearing IncX4 plasmids is associated with variant in replication protein of IncX4 plasmids"

### Supplemental Figures

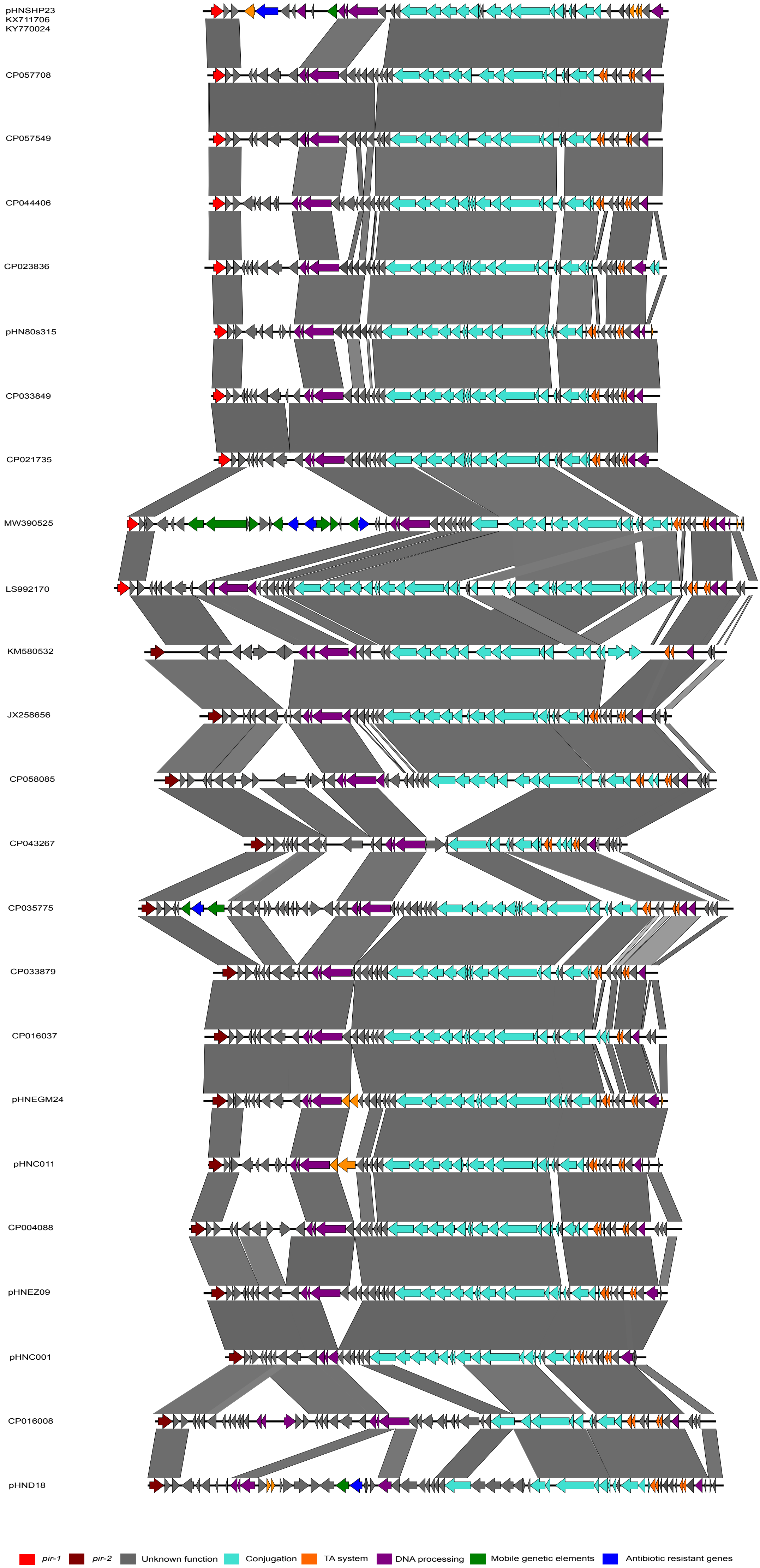

### Supplemental Figures

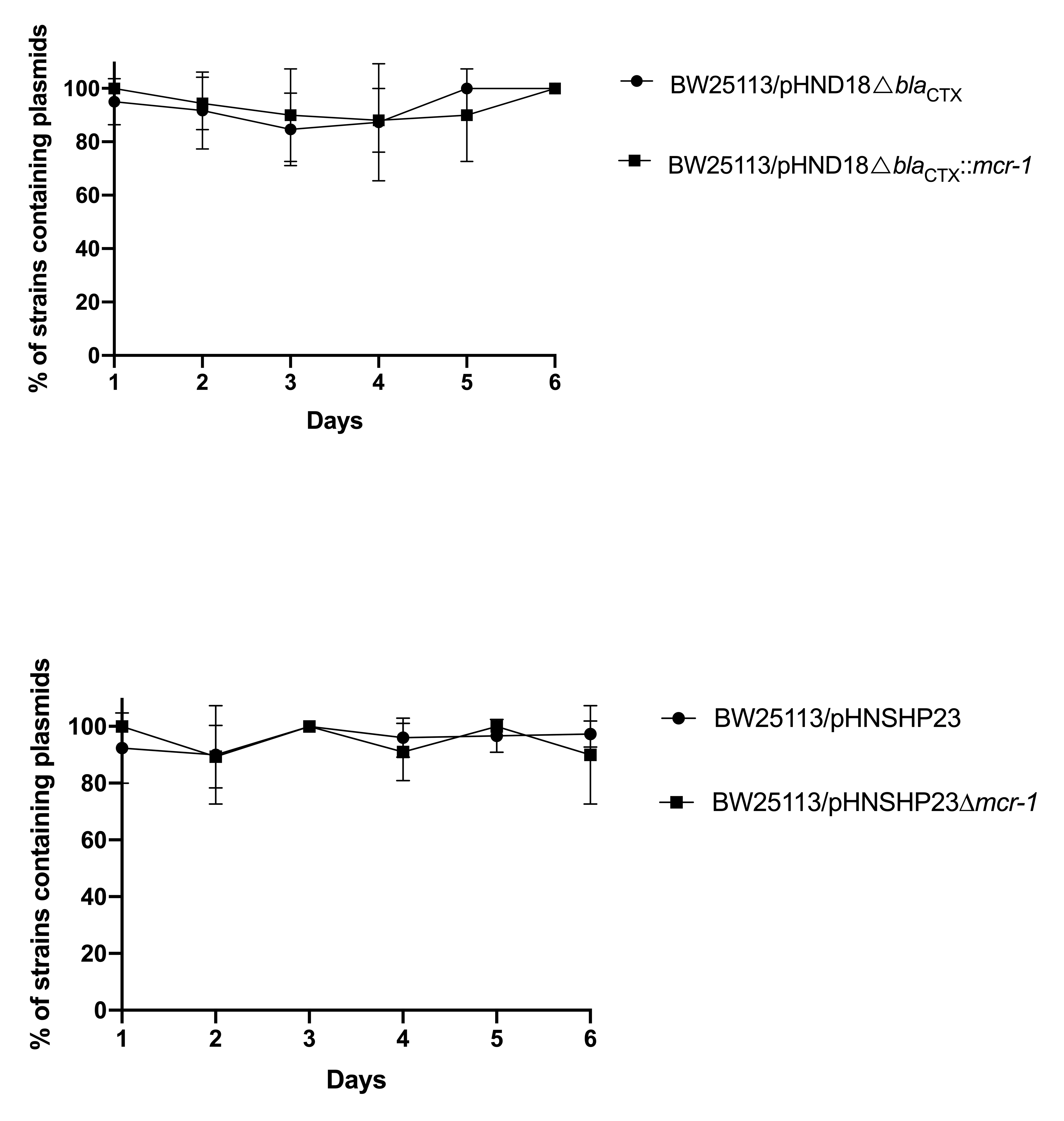
