## Supplemental Figures for "Successful spread of *mcr-1*-bearing IncX4 plasmids is associated with variant in replication protein of IncX4 plasmids"

```
PiR-1     1  ..MRMTTNKTSLSRLTKVRHRNELNSTLSTLPMAAKKVLFLAMCQINSKNE.........
PiR-2     1  MVLKNNKNSDCNDVQSLLAQGNQLLEGAYDITLIEMRLLYLALTKIDSRKPQPANEYTLF

PiR-1    50  ........................FDDDHIFYVTVADYIKWVQVKPDAAYLALRDGSNI
PiR-2    61  AKEYRDTFSLDSKNCYEQLKSAASSLGSKPIVTYEWNEAKKRIDAVKRFWFSSIRYGVGN

PiR-1    85  LDTTLLKLKHDEILELSSDLGFKFTKSNVPDSMNLSLTVFS...TYYRNEGRVGIKFTKE
PiR-2   121  SESDITLRFSDSVSQYLYELKSEFTQMNLEHMVKLDTPFSFRLYSWLYKYKNLNRNKKHS

PiR-1   142  AKRFLCKLIGEEKRYTTQVLLSVVRLTSVNAASLYQLIRKIYSNNSRANSFEMTIDELKD
PiR-2   181  GVISTDPITIEWMKERTGLTGKYPVYKDFKKRVLDPAVDIINANTNLSVTYEGIKSGRKI

PiR-1   202  ELNLYTIGAG.....GVKDYKYPDYPAFKRDVLNKSVKEIMKHTEVKNLSFVVSEKIGRK
PiR-2   241  ESVVFTYLVENETHDGKKIATKPLRPRLPSRPRVIKGSSAEVAWAKKCIDVMKNYLISLR

PiR-1   257  VYKLKFSYTIGYEGDTREDSEFTN.MFDKMYPPEN
PiR-2   301  EYDDKLKLSRPDLQKVKLWHEITGWDFEESFWFSD
```
