## Supplemental Tables for "Successful spread of *mcr-1*-bearing IncX4 plasmids is associated with variant in replication protein of IncX4 plasmids"

**Table S2.** Strains and plasmids used in this study

| Strain or Plasmid | Relevant genotype or phenotype | Reference |
| --- | --- | --- |
| *Escherichia coli* |  |  |
| BW25113 | F^_^Δ(araD-araB)567 ΔlacZ4787(::rrnB-3) λ^_^rph-1 Δ(rhaD-rhaB)568 hsdR514 | (1) |
| VB111 | Nx derivative of MG1655(Nx^R^) | (2) |
| VB112 | Rf derivative of MG1655 (Rf^R^) | (2) |
| BL21 | F-*ompT* *hsdSB*(rB- mB-) gal dcm λ(DE3) Ω PtacUV5::T7 polymerase | Novagen |
| C600 | (Sm^R^) | (3) |
| MFDpir+ | MG1655 RP4-2-Tc::[ΔMu1::*aac(3)IV*-Δ*aphA*-Δ*nic35*-ΔMu2::*zeo*] Δ*dapA*::(*erm-pir*) ΔrecA | (4) |
| GDE8P261 | *E. coli* isolated from swine at slaughter in Guangzhou, China | (5) |
| Plasmid |  |  |
| pBAD30 | ori_p15A_*bla* *araC* P_BAD_ (Ap^R^) | (6) |
| pKD3 | Cm^R^ PCR template for one-step chromosomal gene inactivation | (1) |
| pKD4 | Km^R^, PCR template for one-step chromosomal gene inactivation | (1) |
| pCP20 | Thermo-inducible expression of Flp recombinase (Ts^R^ Ap^R^ Cm^R^) | (7) |
| pKD46 | Ap^R^, λRed recombinase expression | (1) |
| pBAD-*pixR* | pBAD30::*pixR* (Ap^R^) | This study |
| pBAD-*cds9* | pBAD30::*cds9* (Ap^R^) | This study |
| pET28b | Km^R^, lacI^q^ | Novagen |
| pET28b-pilxR | pET28b::*pixR* (Km^R^) | This study |
| pOPlacZ | pAH56 lacZ (Km^R^) | (8) |
| pFG036 | *ori*_pMB1_, *cI857* (Ts^R^) repressor, *tetM* (Tc^R^) | Addgene #137996 |
| pFG051 | *ori*_R6K_, Tn5 *tnp* under λPL promoter, *oriT*_RP4_, Tn5*d-aadA7* (Sp^R^) | Addgene #137997 |
| pHSG575 | *oriV*_pSC101_*(*Cm^R^*)* | (9) |
| pHSG575-*cds9* | pHSG575 carrying *cds9* gene with native promoter | This study |
| pHSG575-*pixR* | pHSG474 carrying *pixR* with native promoter | This study |

Ap, ampicillin; Cm, chloramphenicol; Km, kanamycin; Nx, nalidixic acid; Rf, rifampicin; Sm, streptomycin; Sp, spectinomycin; Tc, tetracycline; Ts, thermosensitive.
